# DEEPReditor-CMG: A deep-learning-based predictive RNA editor for crop mitochondria genomes

**DOI:** 10.1101/2025.10.09.681429

**Authors:** Sidong Qin, Zhihao Xu, Yongqiang Wang, Ziqi Wang, Yixiang Cao, Chaoliang Hou, Jiangshan Yang, Yijie Liu, Yi Xiao, Zhijun Dai, Wei Zhou, Qin Li

**Author notes:** These authors contributed equally to this work (Sidong Qin, Zhihao Xu).

## Abstract

C-to-U RNA editing is a pervasive phenomenon in crop mitochondria, playing a pivotal role in the processing of functional proteins encoded by mitochondrial genes and serving as a crucial regulatory mechanism for nuclear control over mitochondrial gene expression. However, identifying editing sites across the mitochondrial genome with high precision remains a formidable challenge. To address this challenge and uncover the potential regulatory patterns of C-to-U RNA editing, this study compiled a comprehensive dataset of mitochondrial genome sequences from 13 crops, including *Arabidopsis thaliana, Lactuca sativa var. Capitata, Oryza sativa Japonica*, and *Zea mays*, among others, annotated with editing sites from NCBI. Utilizing TensorFlow 2.0, we constructed a rigorous hyperparameter optimization framework, which facilitated the development of the DEEPReditor-CMG model, demonstrating superior predictive capabilities. Based on the DEEPReditor-CMG model and three interspecies relationship indices, we discovered that species with closer phylogenetic relationships exhibit more similar C-to-U RNA editing mechanisms, providing valuable insights into the evolutionary conservation of RNA editing mechanisms. We further developed executable programs for predicting C-to-U RNA editing sites in crop mitochondrial genomes, leveraging the conserved nature of the crop RNA editing machinery to maximize prediction reliability and uncover new editing sites with great potential. This research not only deepens our understanding of C-to-U RNA editing and post-transcriptional regulatory mechanisms in crop mitochondria but also paves a novel avenue for investigating RNA editing mechanisms. Experimental data, computational code, and applications are publicly accessible at https://github.com/Qinsidong/DEEPReditor-CMG.

**Highlights:** High-precision prediction model for C-to-U RNA editing in crops—DEEPReditor-CMG

Three interspecies relationship indices describing C to U RNA editing

Prediction tools based on the conservation of crop RNA editing mechanisms

Open-sourced experimental data, computational code, and applications

## 1. Introduction

RNA editing is a phenomenon characterized by the targeted alteration of codons through nucleotide modifications at the RNA level, which can be accomplished via substitution, insertion, or deletion of nucleotides (Brennicke, Marchfelder & Binder, 1999). This process is essential for the generation of functional proteins by mitochondrial genes and serves as a significant regulatory mechanism for nuclear control over mitochondrial gene expression (Volloch, Schweitzer & Rits, 1990; Mulligan, Williams & Shanahan, 1990). Since its initial discovery in the 20th century, the molecular mechanisms underlying RNA editing have been the subject of extensive research (Hiesel, 1989; He et al., 2018; Small et al., 2020; Zheng et al., 2020; Edera et al., 2021; Wang et al., 2021, Ali et al., 2025).

Among the various forms of RNA editing, C-to-U RNA editing is the most prevalent in crop species (Simpson & Emeson, 1996). The identification of C-to-U RNA editing sites presents a considerable challenge, requiring methods that are efficient, rapid, and cost-effective. Traditional experimental workflows for the identification of RNA editing sites are complex, multi-step, and costly. While these methods typically yield accurate results, they are often limited to the identification of editing sites for single genes within the coding regions. However, it is now recognized that RNA editing in non-coding regions can also exert regulatory effects on gene expression (Nishikura, 2016; Zhang, 2017). Furthermore, the entire experimental process is susceptible to various sources of interference, such as PCR errors, the confounding effects of DNA editing at SNPs and mutants, and the influence of cellular heterogeneity on gene expression levels (Grohmann et al., 2016; Lerner et al., 2021). These factors necessitate the employment of multiple technical strategies to mitigate the impact of erroneous results during the identification process.

The proliferation of machine learning research has opened up new frontiers for the high-precision prediction of RNA editing sites. Previous endeavors to predict RNA editing sites using conventional machine learning techniques, such as decision trees, random forests, and support vector machines (SVM), have been pursued by numerous scholars (Kim, Hur & Kim, 2016; Sun et al., 2016; Chen et al., 2017; Cummings & Myers, 2004; Mower, 2005; Thompson & Gopal, 2006; Du, He & Li, 2007; Lenz & Knoop, 2013). However, these methods typically yield suboptimal predictive accuracy and often overlook the interspecies variations that could significantly influence the editing process. In a previous study in 2022, we introduced a novel modeling approach that leverages multiple feature extraction to highlight the potential for greater mechanistic similarities in C-to-U RNA editing among closely related species (Qin et al., 2022). This insight not only enhances our understanding of the evolutionary conservation of RNA editing mechanisms but also paves the way for innovative cross-species prediction models. The findings underscore the importance of considering phylogenetic relationships in the development of predictive algorithms for RNA editing, offering a promising avenue for future research and applications in the field of comparative genomics and RNA biology.

The manually set feature extraction method significantly impacts the accuracy of traditional machine learning models (Fang et al., 2022). While these features are often too abstract and pose challenges for interpretation, deep learning’s autonomous learning capabilities can circumvent human cognitive limitations and more accurately capture underlying patterns, thereby improving model fit (Hinton & Salakhutdinov, 2006). Rumelhart et al. (1986) introduced the Back Propagation Network (BPN), which learns representations by back-propagating errors and has since garnered considerable attention in deep learning research. Among deep learning models, Convolutional Neural Networks (CNNs) are notable for their local connections, weight sharing, pooling operations, and multi-layer structures (Lecun, 2015). These features endow CNNs with powerful automatic feature learning capabilities, making them widely applicable in image and text recognition fields (He et al., 2020; Bazgir, Ghosh & Pal, 2021; Bertoni, Citti & Sarti, 2022, Han et al., 2024).

Genomic data, at its core, consists of textual sequences of base pairs, rendering the CNN structure an ideal framework for our model. This study meticulously constructed a stringent hyperparameter optimization process based on the crop mitochondrial genomes and fragments, culminating in the development of a highly accurate prediction model dubbed DEEPReditor-CMG (a deep learning-based predictive RNA editor for crop mitochondria genome). Beyond mere prediction, this research delved into the interspecies correlation of C-to-U RNA editing mechanisms, employing a suite of interrelationship evaluation metrics. In a final step, executable programs were developed to predict C-to-U RNA editing sites across the entire genome and within specific fragments. All python project processes were open-sourced to facilitate collaborative advancement in the field. This study marks a significant stride in the predictive modeling of RNA editing events in crop mitochondria, offering a valuable tool for future research and applications in plant genomics.

## 2. Materials and methods

### 2.1 Collection of materials

In this study, we amassed a comprehensive collection of whole mitochondrial genomes from a diverse array of crop species. These genomic datasets were meticulously downloaded from the National Center for Biotechnology Information (NCBI), which stands as a reputable repository for biological data. Alongside the genomic sequences, we also procured the corresponding annotations for RNA editing sites. These annotations are instrumental in identifying the precise locations within the genome that undergo C-to-U RNA editing, providing a foundational dataset for our subsequent analyses.

### 2.2 Composition of positive and negative samples

According to the mitochondrial genome sequences and their respective RNA editing site annotations, positive samples were composed centered on editing site C, whereas negative samples were composed centered on C lacking editing site annotations. Each sample was flanked by a 250 bp extension both upstream and downstream from the central C, allowing for the inclusion of potential regulatory elements and sequence motifs that may influence the editing process (Du He & Li, 2007; Qin et al., 2022). To maintain dataset balance and avoid biases towards positive samples, negative samples were extracted randomly and in equal numbers to the positive samples (see supplementary material “ExtractSamples.py” for sample extraction code:

https://github.com/Qinsidong/DEEPReditor-CMG/blob/main/ExtractSamples.py).

### 2.3 Training, validation, and test set partitioning

To ensure the generalizability and robustness of our model, the dataset was meticulously partitioned into a training set and an independent test set in an 8:2 ratio. A portion of the training set, specifically 20%, was further allocated as the validation set to monitor the model’s performance during the training process and to prevent overfitting. This stratified random sampling was conducted five times, employing varying ‘random states’ to ensure that the dataset divisions were not biased towards any particular subset of data (see supplementary material “ExtractSamples.py”). Within the training dataset, we maintained a balanced proportion of positive and negative samples. This equilibrium is essential for mitigating the risk of sub-optimal parameter estimation and potential distortions in model evaluation. (Krawczyk, 2016).

### 2.4 Model architecture

Our CNN model was meticulously constructed utilizing Tensorflow 2.0 (https://github.com/tensorflow/tensorflow/releases/tag/v2.0.0) and keras (https://github.com/keras-team/keras), and the model structure was Input layer + Embedding layer + two Convolutional and Pooling layers + Dropout layer (30%) + Sigmoid output layer, batch_size = 50, optimizer was Adam, loss function was “binary_crossentropy” (Fig. 1F), and the evaluation metric was ACC (Equation 1).

**Figure 1.**
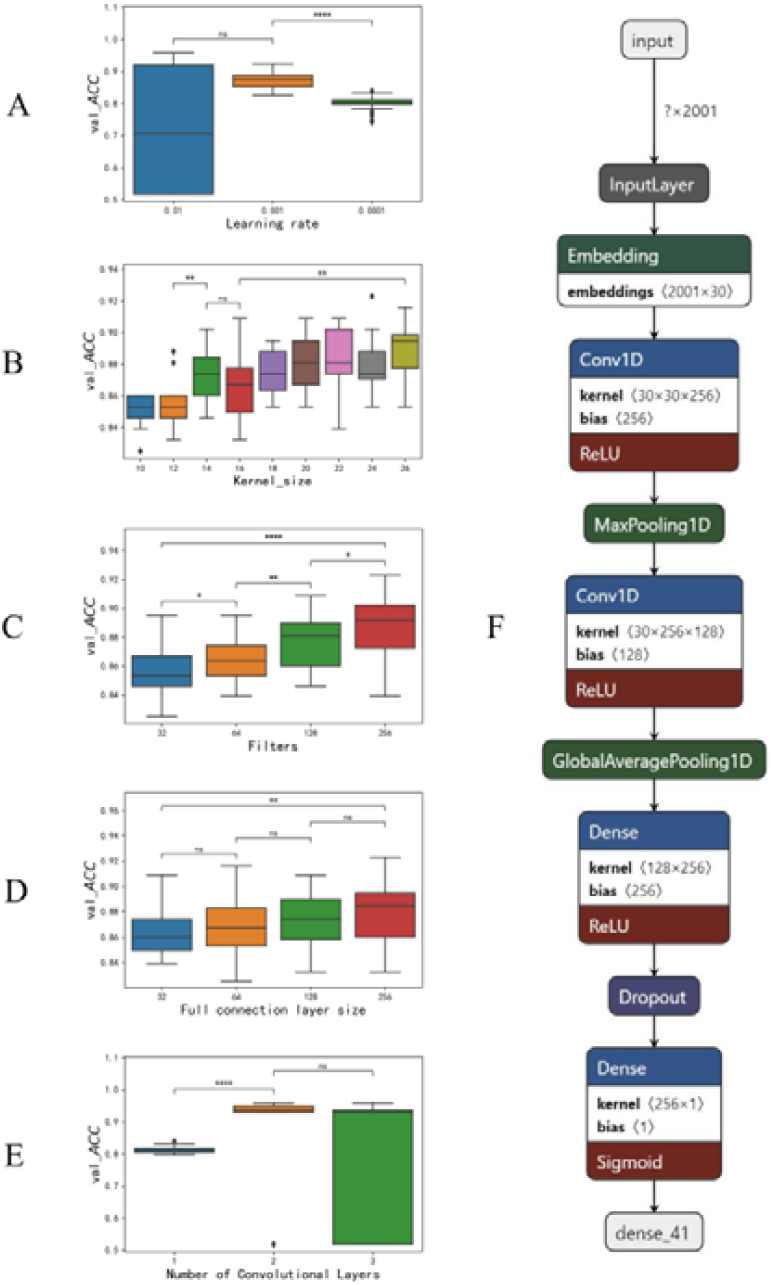
Model hyperparameter range and architecture

### 2.5 Hyperparameter optimization strategy

The optimization of the hyperparameter range in this study was conducted through a synergistic approach combining preliminary screening and range search. This hybrid methodology leveraged the strengths of both random search and grid search techniques, optimizing the balance between exploration and exploitation of the hyperparameter space. Initially, the random search phase was employed to broadly explore the hyperparameter space, identifying promising regions that could potentially lead to improved model performance. Subsequent to the preliminary screening, a grid search was applied to fine-tune the hyperparameters within the identified ranges. This two-stage optimization process enabled a rapid convergence on the most effective hyperparameter settings, significantly reducing computational costs and saving a significant amount of time, so this approach was particularly advantageous for training models on portable machines with limited computational resources. In contrast to some existing studies that have neglected explicit hyperparameter optimization or have failed to provide detailed descriptions of their optimization processes, our study placed a strong emphasis on this critical aspect of model development. By incorporating a rigorous hyperparameter optimization strategy into our experimental design, we ensured that the resulting model was robust and generalizable, thereby enhancing the credibility and reproducibility of our findings.

### 2.6 Interspecific relationship indicators

#### 2.6.1 Interspecies average nucleotide identity (ANI)

ANI is a computational algorithm designed to approximate the DNA-DNA hybridization (DDH) technique at the nucleic acid level. It assesses the similarity between two genomes by calculating the identity of all orthologous protein-coding genes, thereby providing a measure of genetic relatedness (Goris et al., 2007). The Basic Local Alignment Search Tool (Blast) serves as the predominant method for computing ANI due to its expeditious and highly accurate sequence alignment capabilities (Altschul et al., 1990; Lu, Noble & Keich, 2024). In the present study, complete mitochondrial genomes of different species were analysed for ANI and then interspecies ANI values were derived from pairwise comparisons of these species by BLAST method. These values facilitate a detailed examination of the genetic distances and relationships among the selected species, offering valuable insights into the conservation and divergence of mitochondrial genomes across different crops. This approach allows for a robust quantification of interspecies similarities, which is essential for understanding the broader evolutionary context of RNA editing mechanisms in crop mitochondria.

#### 2.6.2 Interspecies prediction analysis

Within the scope of model development, we employed an optimal CNN model for each of all species under investigation. Subsequently, we conducted interspecies prediction analyses to assess the predictive capacity of each model across different species. Specifically, the model trained on species A was utilized to predict editing sites in species B, denoted as “interspecies prediction +”. Conversely, the model trained on species B was employed to predict editing sites in species A, denoted as “interspecies prediction -”. This reciprocal prediction approach was systematically applied to all pairs across the full range of species, thus assessing the model’s ability to generalize across species and providing insights into the conservation of RNA editing patterns among different crops.

#### 2.6.3 Model Euclidean distance assessment

Within the interspecies prediction phase, we modified the architecture of the optimal CNN model for each species. Specifically, we eliminated the last sigmoid layer and then repurposed the penultimate layer with the output layer. This alteration enabled each model to generate a 256-dimensional vector output that captured the nuanced differences between the individual models’ representations.

Subsequently, the positive and negative samples of each species were utilized as inputs to their respective models, a process that would resulted in the derivation of vectors of these positive and negative samples for each species. To ensure the stability of these vectors and to minimize errors, the input to each model consisted of all the samples from a given species and the mean of the vector matrix for each species was calculated.

Finally, the Euclidean distances between the positive and negative sample vectors were computed for each species (Equation 2), yielding a comprehensive set of Euclidean distance measurements. These distances provide a quantitative measure of the dissimilarity between the positive and negative sample representations across species, offering insights into the variability of RNA editing site patterns and the generalizability of our models across different biological contexts.

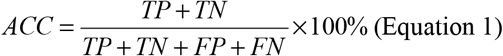

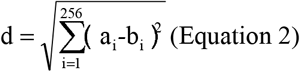

Notes*TP*: true positive, *FN*: false negative, *TN*: true negative, *FP*: false positive; d: Euclidean distance, a_i_: the i-th position of vector a, b_i_: the i-th position of vector b.

## 3. Results and analysis

### 3.1 Dataset collection and analysis

At the outset of our investigation, we amassed a comprehensive collection of complete mitochondrial genomes from 13 diverse crop species, complemented by their respective RNA editing site annotations, retrieved from the NCBI (Table 1). This curated dataset spans a broad taxonomic range, offering a panoramic view of the RNA editing landscape within crop mitochondria. The compilation of these genomic resources has enabled a detailed exploration of the RNA editing patterns and has set the stage for in-depth analyses of the underlying mechanisms governing these post-transcriptional modifications in selected crop species.

**Table 1.**
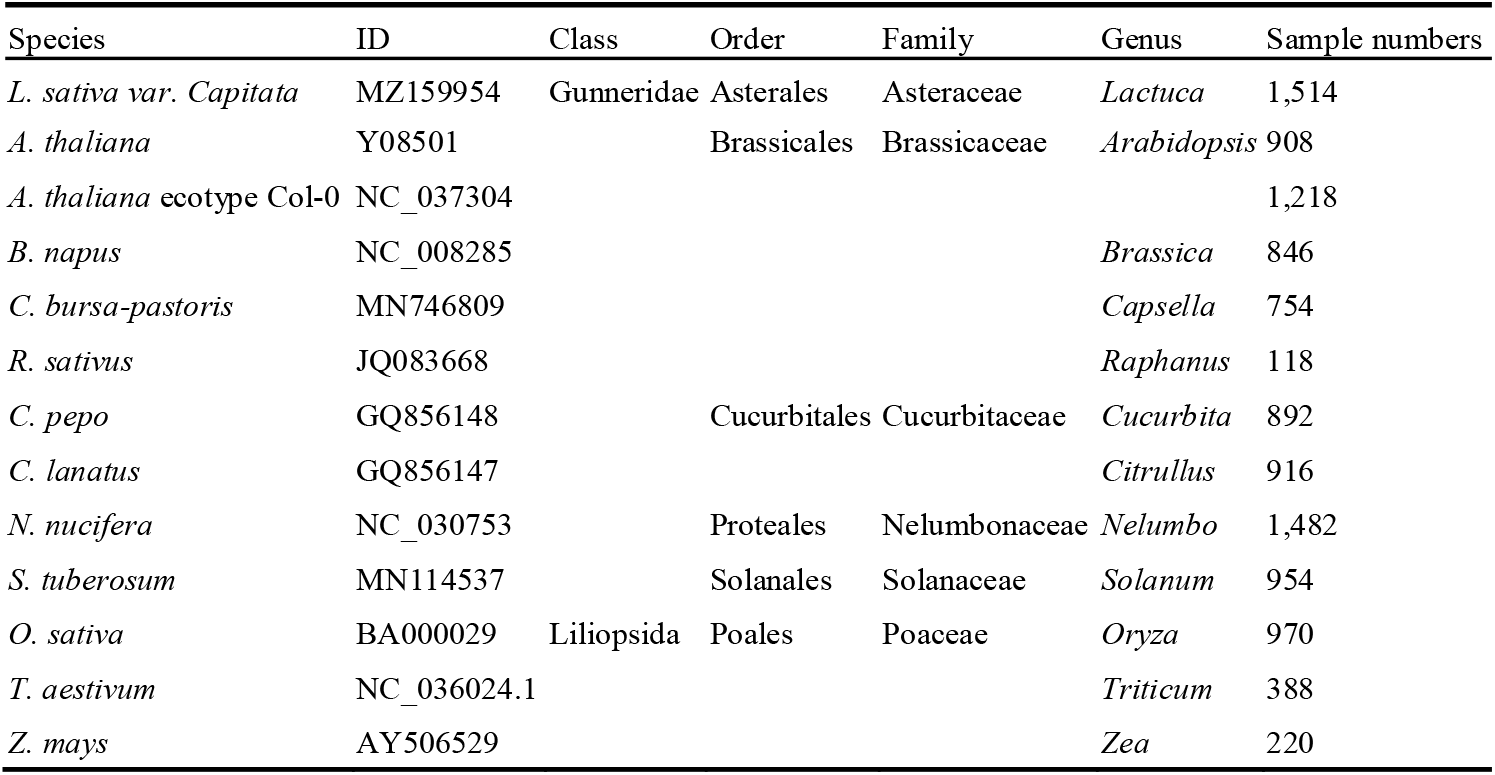
Taxonomic details and accession numbers of the thirteen crops.

A noteworthy exception was the mitochondrial reference genome of *Triticum aestivum* (NC_036024.1), which, despite lacking direct annotation of editing sites, offered a valuable resource in the form of 11 mitochondrial genome fragments (X57965, X57968, Y13920, AB022060, AJ295996, X69720, X75036, X79609, X69205, AF082025, and AJ535507). By meticulously analyzing these fragments, we were able to infer the editing site information for *T. aestivum* indirectly. In contrast, for the remaining species, the editing site annotations were obtained directly from their respective whole mitochondrial genome annotations.

### 3.2 DEEPReditor-CMG construction

In this study, we randomly selected *Cucurbita pepo* as a representative species to evaluate the impact of various hyperparameters on the performance of single-layer convolutional network models. We established models with learning rates of 0.1,0.001 and 0.0001 and systematically varied the “Kernel_size” of the convolutional layer from 10 to 26 in steps of 2, with the embedding layer adjustments synchronized accordingly; the “Filters” were scaled from 32 to 256 in doubling increments, and the “Full connection layer size” was similarly adjusted, resulting in a total of 144 single-layer convolutional models for each learning rate.

Our analysis revealed that at a learning rate of 0.01, 74 out of 144 models failed to achieve a satisfactory fit, with a validation accuracy (val_ACC) of 0.5. In contrast, models with learning rates of 0.001 and 0.0001 exhibited better fit and stability, with the 0.001 rate performing significantly better than the 0.0001 rate (Fig.1A). The influence of “Kernel_size” on the validation set accuracy was subtle but suggested a positive correlation; model performance was poorer when “Kernel_size” was less than 14 and peaked at an optimal accuracy when “Kernel_size” was 26 (Fig.1B). An increase in “Filters” corresponded to higher validation set accuracy (Fig.1C), while the “Full connection layer size” had a minimal impact on accuracy, with the best results obtained when set to 256 (Fig.1D).

In accordance with the empirical findings from single-layer convolutional neural networks, we set the learning rate to 0.001 for further optimization. For each layer, the “Kernel_size” was systematically varied from 10 to 26 with an increment of 2. The “Filters” configuration was as follows: the first layer was set to 256, the second layer alternated between 128 and 256, and the third layer combined 64 and 256. The “Full connection layer size” was uniformly set to 256 across all models. Consequently, we developed three distinct convolutional neural network models, each with 18 unique parameter configurations (Fig.1E). Upon evaluation, we observed that two and seven models in the two-layer and three-layer convolutional networks, respectively, failed to achieve a satisfactory fit, exhibiting a validation set accuracy of 0.5. Comparative analysis revealed that the two-layer convolutional structure not only outperformed the single-layer convolutional structure in terms of validation set accuracy but also demonstrated superior stability compared to the three-layer convolutional structure. These observations indicate that the complexity introduced by additional layers can lead to overfitting or underfitting, depending on the specific configuration. The two-layer structure strikes a balance between the model’s capacity to learn from the data and its generalization ability to new, unseen data. Therefore, the two-layer convolutional network was deemed the most effective architecture for our purposes, offering a combination of high predictive accuracy and model stability.

Ultimately, the two-layer convolutional network structure was selected as the final model architecture (Fig.1F), with “kernel_size” ranging from 16 to 30 in steps of 2 for each layer, “Filters” set to 128 and 256, “Full connection layer size” to 256, learning rate to 0.001, optimizer as Adam, loss function as “binary_cross_entropy”, and the evaluation metric as ACC, with the optimal number of iterations dynamically selected. This model architecture was chosen for its balance of accuracy and stability, offering a robust framework for predicting RNA editing sites in crop mitochondria.

### 3.3 Model intraspecific assessment

In our comparative analysis with existing methods, our DEEPReditor-CMG demonstrated superior predictive accuracy in three reference species (*Arabidopsis thaliana, Brassica napus* and *Oryza sativa* Japonica), with 13.41%, 6.90% and 14.46% improvements in test accuracy (test_ACC) compared to the PREP-Mt method, 9.41%, 9.41% and 14.46% compared to the Du-SVM method, 2.20%, 10.71% and 17.28% compared to the iPReditor-CMG method, respectively (Table 2). These results underscore the robustness and efficacy of our novel approach in predicting C-to-U RNA editing sites.

**Table 2.** Comparative analysis of *ACC* between DEEPReditor-CMG and reference prediction methods.

| Species | PREP-Mt | Du-SVM | iPReditor-CMG | DEEPReditor-CMG |
| --- | --- | --- | --- | --- |
| *A. thaliana* | 0.82 | 0.85 | 0.91 | **0.92±0.01** |
| *B. napus* | 0.87 | 0.85 | 0.84 | **0.91±0.02** |
| *O. sativa* | 0.83 | 0.83 | 0.81 | **0.91±0.04** |

Note: In the DEEPReditor-CMG approach, the training and independent test sets were randomly partitioned on five separate occasions employing different random states for each division to ensure variability and robustness in the dataset distribution. The font of the optimal results in the table is in bold.

In contrast to the iPReditor-CMG method, which employed a 10% independent test set, our DEEPReditor-CMG enhanced the validation process by utilizing a 20% independent test set. This increment in the test set size, coupled with the introduction of five distinct randomizations to partition the test set, ensures a more robust and comprehensive assessment of model performance. DEEPReditor-CMG not only amplified the representativeness of the test data but also fortified the reliability of the results by mitigating the impact of any potential bias inherent in a single randomization.

Moreover, within the 13 species, our auto-optimized range hyperparameter model outperformed the independent prediction results of the BPN (“KerasTunerBPN.py”: https://github.com/Qinsidong/DEEPReditor-CMG/blob/main/Hyperparametric_Optimization_Search/Final_model_structure/KerasTunerBPN.py) and the Empirical Optimization CNN (an empirical artificial hyperparameter optimization model) (Table 3). This comparative analysis not only validates the necessity and effectiveness of our range hyperparameter optimization but also suggests that our method is better equipped to capture the underlying coding mechanism of C-to-U RNA editing across diverse species. The superior performance of DEEPReditor-CMG across multiple species and datasets confirms its potential as a powerful tool in the field of RNA editing research.

**Table 3.** *ACC* results of four methods across diverse species.

| Species | BPN | SSBPN | EHCNN | DEEPReditor-CMG |
| --- | --- | --- | --- | --- |
| *L. sativa* var. *capitata* | 0.65 | 0.65 | **0.93** | **0.93±0.01** |
| *A. thaliana* | 0.65 | 0.66 | 0.84 | **0.92±0.01** |
| *A. thaliana* ecotype Col-0 | 0.64 | 0.64 | 0.89 | **0.92±0.01** |
| *B. napus* | 0.61 | 0.66 | 0.86 | **0.91±0.02** |
| *C. bursa-pastoris* | 0.57 | 0.64 | 0.89 | **0.90±0.02** |
| *R. sativus* | 0.65 | 0.81 | 0.87 | **0.91±0.02** |
| *C. pepo* | 0.57 | 0.66 | 0.82 | **0.94±0.02** |
| *C. lanatus* | 0.61 | 0.7 | **0.91** | **0.91±0.02** |
| *N. nucifera* | 0.64 | 0.7 | 0.91 | **0.93±0.01** |
| *S. tuberosum* | 0.62 | 0.66 | 0.87 | **0.89±0.03** |
| *O. sativa* | 0.55 | 0.64 | 0.83 | **0.91±0.04** |
| *T. aestivum* | 0.60 | 0.66 | 0.83 | **0.84±0.06** |
| *Z. mays* | 0.57 | 0.51 | 0.86 | **0.93±0.02** |

Note: In the DEEPReditor-CMG approach, the training and independent test sets were randomly partitioned on five separate occasions employing different random states for each division to ensure variability and robustness in the dataset distribution. The font of the optimal results in the table is in bold.

### 3.4 Interspecies relationships in the C-to-U RNA editing machinery

We conducted an ANI analysis on the complete mitochondrial genomes of 13 species, with *T. aestivum* serving as the mitochondrial reference genome. Among these species, which included 10 dicotyledonous and 3 monocotyledonous plants, we observed a significant negative correlation between “interspecies ANI” and “model Euclidean distance” for both edited (*R*=-0.31, *p*=0.006) and unedited (*R*=-0.35, *p*=0.002) states (Fig. 2). This correlation was significantly strengthened within the dicotyledonous plants, with edited (*R*=-0.66, *p*=9.68e-7) and unedited (*R*=-0.64, *p*=2.74e-6) states showing a more pronounced negative correlation (Fig. 3). However, due to the limited number of monocotyledonous plants, the strong negative correlation observed in this group, while present, is a reliability that needs to be supported by more genomes of the species to be tested. Our findings suggest that the smaller the “model Euclidean distance” between two species, the more similar their C-to-U RNA editing mechanisms, indicating a closer phylogenetic relationship. In contrast, species that are not closely related exhibit greater differences in their C-to-U RNA editing mechanisms.

**Figure 2.**
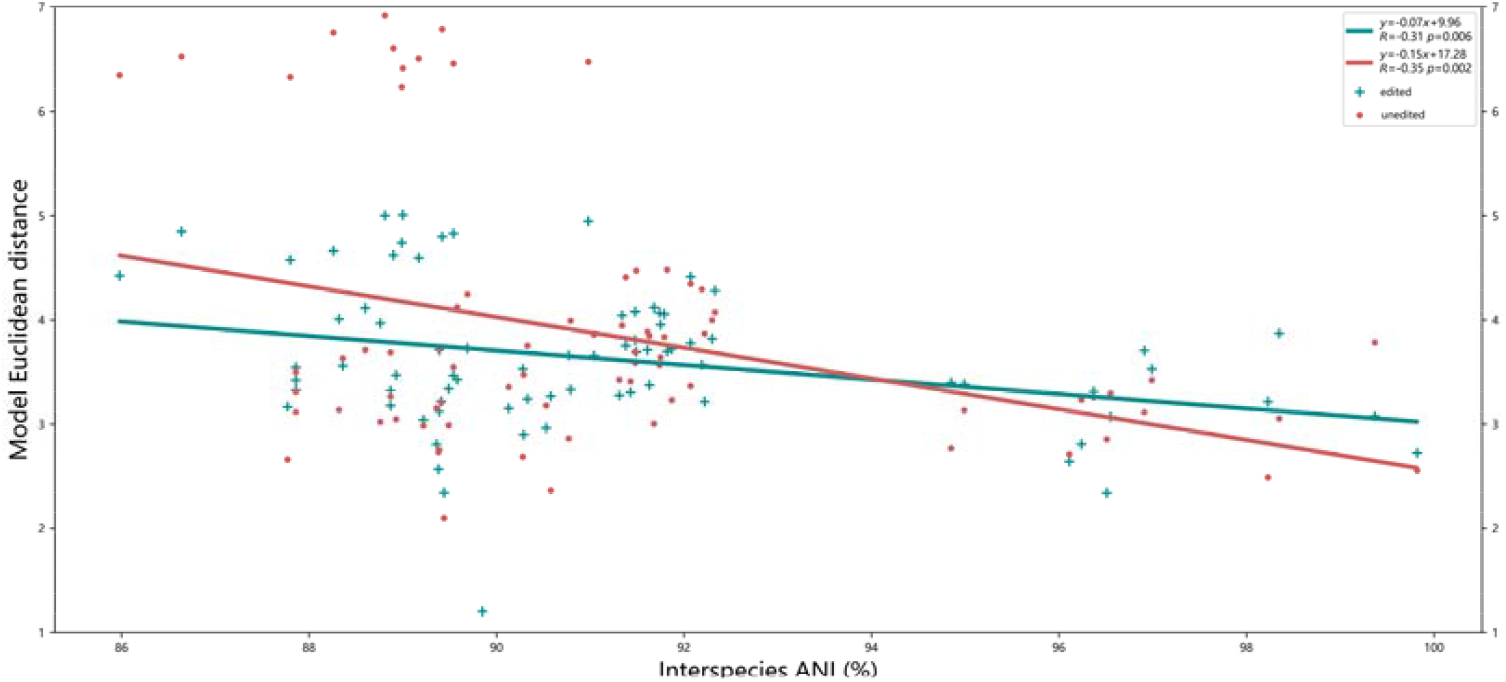
Relationships of “interspecies ANI” and “model Euclidean distance” among 13 species

**Figure 3.**
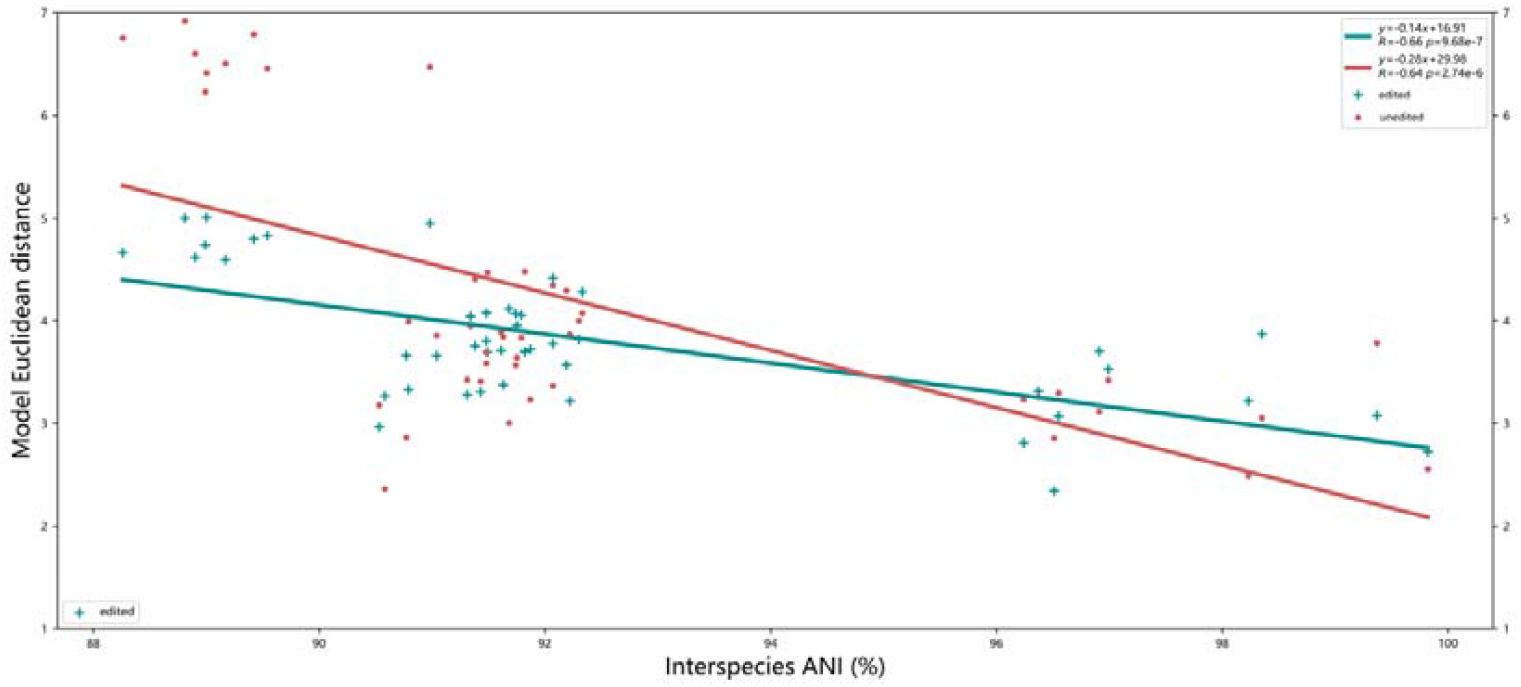
Relationships of “interspecies ANI” and “model Euclidean distance” in dicotyledonous plants

Considering the directional nature of “interspecies prediction” values, which fundamentally indicate the similarity of RNA editing mechanisms between species, we chose the maximum value for “interspecies prediction” between pairs of species. A significant positive correlation (*R*=0.45, *p*=4e-5) was found between “interspecies ANI” and “interspecies prediction” across the 13 species (Fig. 4), aligning with the conclusions drawn from “model Euclidean distance”: the more phylogenetically close two species are, the more similar their RNA editing mechanisms.

**Figure 4.**
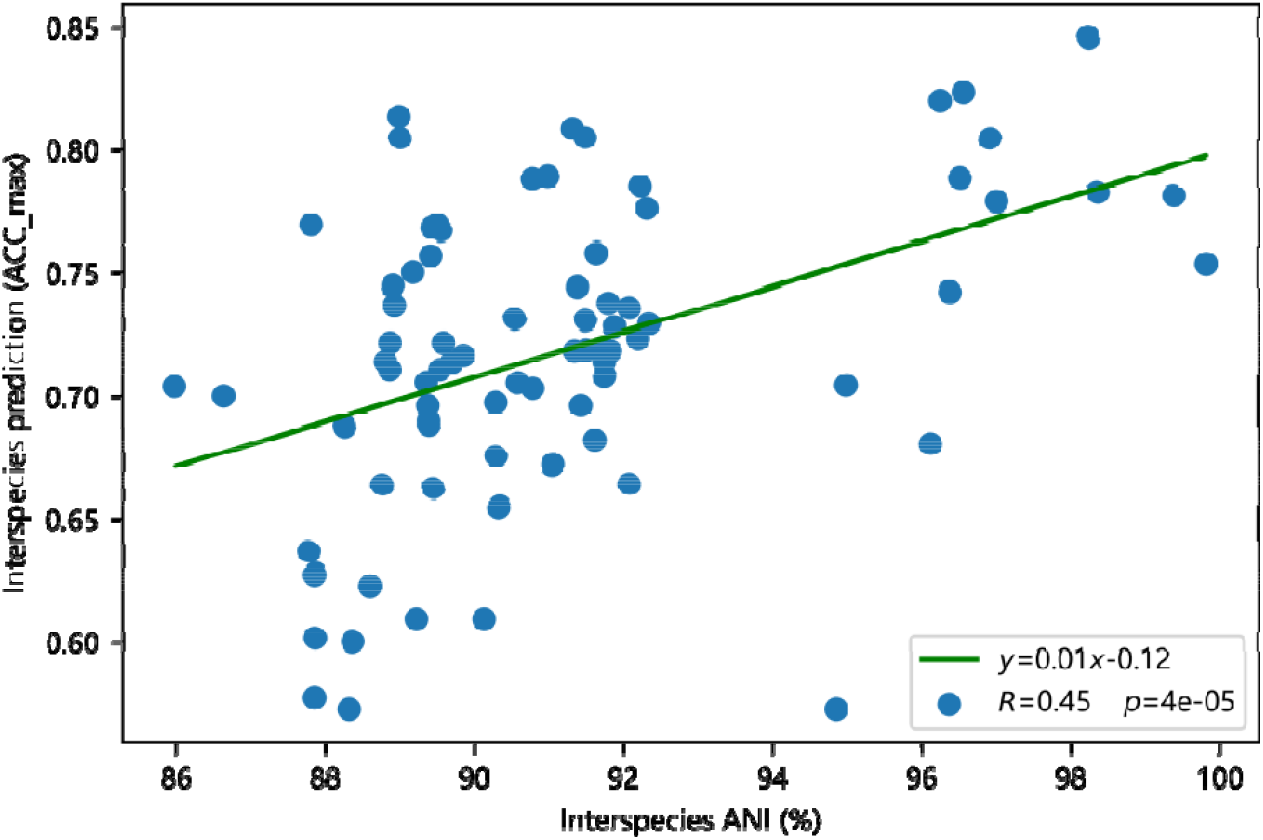
Relationships of “interspecies ANI” and “interspecies prediction” among 13 species

“Interspecies ANI” reflects the phylogenetic relationship between species at the genomic level, while “model Euclidean distance” and “interspecies prediction” based on high-precision deep learning (DL) models, indicate the similarity of C-to-U RNA editing patterns and mechanisms among different species. We found “interspecies ANI” to be negatively correlated with “model Euclidean distance” and positively correlated with “interspecies prediction” (all detailed evaluation values were presented in Supplementary Table 1). These correlations further support the inference made by our previous iPReditor-CMG method, demonstrating that phylogenetically closer species have more similar C-to-U RNA editing mechanisms. Therefore, when modeling and predicting, it is crucial to consider the phylogenetic relationships among different species rather than merely combining datasets from different species. This significant conclusion not only provides theoretical support for the application of our study but also underscores the importance of phylogenetic context in the interpretation of RNA editing mechanisms.

### 3.5 Practical application and utility for DEEPReditor-CMG

#### 3.5.1 Softwar optimization and development based on phylogenetic proximity

This research revealed a heightened similarity in RNA editing mechanisms among closely related species. In alignment with this finding, we optimized and evaluated the selection criteria for species prediction models to enhance the utility of the DEEPReditor-CMG tool. The independent test results suggested a preference for high-precision models constructed from corresponding species when the species to be predicted were among the 13 species examined in our study. For predictions extending beyond these 13 species, a preference was given to models derived from species sharing the same genus, order, or class, based on their taxonomic affinities (Table 4). Our DEEPReditor-CMG ensures that the DEEPReditor-CMG tool leverages the most relevant and accurate models for predicting RNA editing sites, thereby improving the reliability of its predictions and tailoring its functionality to the specific needs of diverse species.

**Table 4.** *ACC* results of the amplification models.

| Genus | Test_ACC | Family | Test_ACC | Class | Test_ACC |
| --- | --- | --- | --- | --- | --- |
| *Arabidopsis* | 0.92 | Brassicaceae | 0.94 | Gunneridae | 0.94 |
|  |  | Cucurbitaceae | 0.92 | Liliopsida | 0.87 |

The DEEPReditor-CMG is equipped to predict user-specified samples using the script “PreCustomData.py” (https://github.com/Qinsidong/DEEPReditor-CMG/blob/main/DEEPReditor(Python_scripts_version)/PreCustomData.py) and to conduct whole-genome predictions with “PreWholeGen.py” (https://github.com/Qinsidong/DEEPReditor-CMG/blob/main/DEEPReditor(Python_scripts_version)/PreWholeGen.py). Additionally, executable programs have been developed for convenient local use by end-users, providing a flexible tool for a variety of genomic prediction tasks.

#### 3.5.2 Genome-wide prediction analysis

Taking *Cucurbita pepo* (ID: GQ856148), the mitochondrial genome upon which the DEEPReditor-CMG model was trained, as a case study to assess the genome-wide prediction of RNA editing sites. Given the vastness of the whole genome, the annotation of editing sites was less than one-thousandth, making this analysis a critical test for the reliability of the DEEPReditor-CMG method and a valuable opportunity to uncover potential editing sites in this species. The DEEPReditor-CMG platform demonstrated its effectiveness by successfully identifying 18 out of the 22 known editing regions, as indicated by red boxes in Figure 5, with an accuracy rate exceeding 80% at the mitochondrial genome-wide level. Moreover, the platform strongly indicated the potential presence of previously unidentified editing sites within the 0.7e6 to 0.8e6 bp region of the mitochondrial genome, so the finding provide actionable insights for future experimental validation.

**Figure 5.**
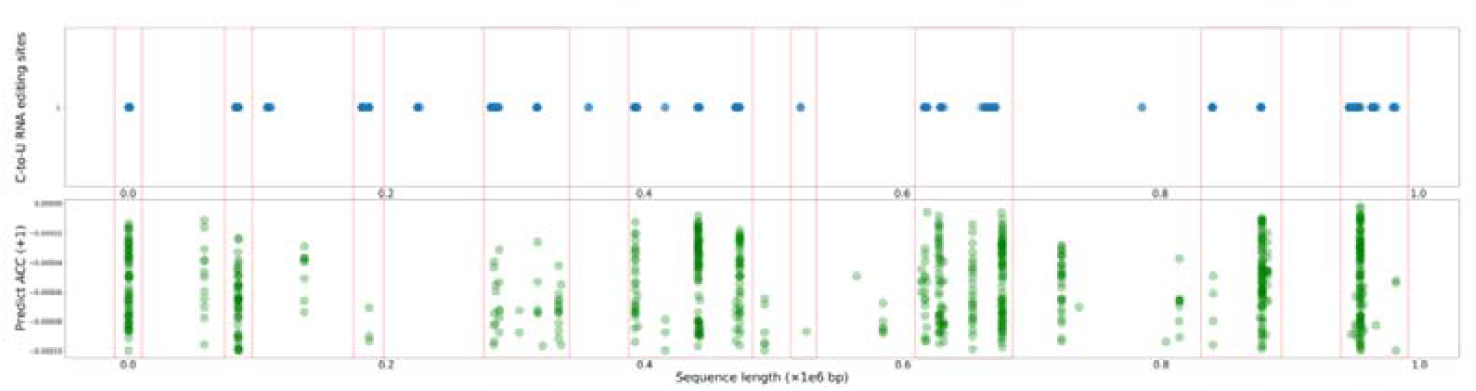
Mitochondrial genome of *Cucurbita pepo*: correlation between predicted editing sites and NCBI annotations

These results not only underscore the efficacy of our approach in predicting editing sites across the crop mitochondrial genome but also highlight its significant potential to inform and guide experimental research. The DEEPReditor-CMG platform’s performance in this case study validates its utility as a predictive tool and emphasizes its role in advancing our understanding of RNA editing mechanisms in crop species

Note: Blue dots denoted C-to-U RNA editing sites annotated by NCBI, whereas green dots represented the DEEPReditor-CMG predicted editing sites. The intensity of the color indicates the density of sites in the proximity of that region.

## 4. Discussion

The field of RNA editing mechanisms and prediction of editing sites through machine learning is burgeoning, with a plethora of studies contributing to the understanding of this complex post-transcriptional modification (Cummings & Myers, 2004; Mower, 2005; Thompson & Gopal, 2006; Du, He & Li, 2007; Lenz & Knoop, 2013). While the iPReditor-CMG method (Qin et al., 2022) has achieved commendable prediction accuracy in models based on three crops and has begun to explored inter-species relationships in RNA editing mechanisms, it is constrained by a limited number of species. The feature extraction-based model construction in previous methods has proven to be inadequate for capturing the nuances of RNA editing mechanisms, particularly when it comes to cross-species mitochondrial genome-wide prediction, which remain a formidable challenge.

Our novel DEEPReditor-CMG, grounded in the mitochondrial genomes of 13 diverse crops, harnesses the robust autonomous feature learning capabilities of CNN. Through a rigorous parameter optimization process, we have trained a high-precision prediction model. The accuracy of our independent test set not only surpasses previous methods in the reference species but also demonstrates robust performance across all 13 crops. Utilizing three inter-species relationship evaluation indicators interspecies ANI, interspecies prediction, and model Euclidean distance—we have corroborated through correlation analysis that the closer the phylogenetic proximity of two species, the more similar their C-to-U RNA editing mechanisms are. This finding not only reinforces the hypothesis proposed by the iPReditor-CMG method, but also paves new avenues for interspecific prediction. Furthermore, our DEEPReditor-CMG’s ability to bring the model closest to the taxonomic relationships of the species of interest through incremental matching based on genus, family, order, and class contributes to improved prediction reliability. More notably, DEEPReditor-CMG is not limited to bulk predictions of editing sites at specified locations, but also predicts editing sites across the entire mitochondrial genome of a species, providing strong support for mining potential sites.

To facilitate the application of our method, we have made our source code publicly available and developed an executable program for enhanced local usability (https://github.com/Qinsidong/DEEPReditor-CMG). This research not only provides a preliminary analysis of experimental data for related researchers, potentially narrowing the verification scope and reducing experimental costs, but also offers new research avenues for a deeper comprehension of RNA editing mechanisms. Looking ahead, we intend to expand the range of species, ensuring that the powerful predictive capabilities of DEEPReditor-CMG extend beyond crop mitochondrial genomes, serving researchers across various disciplines.

---

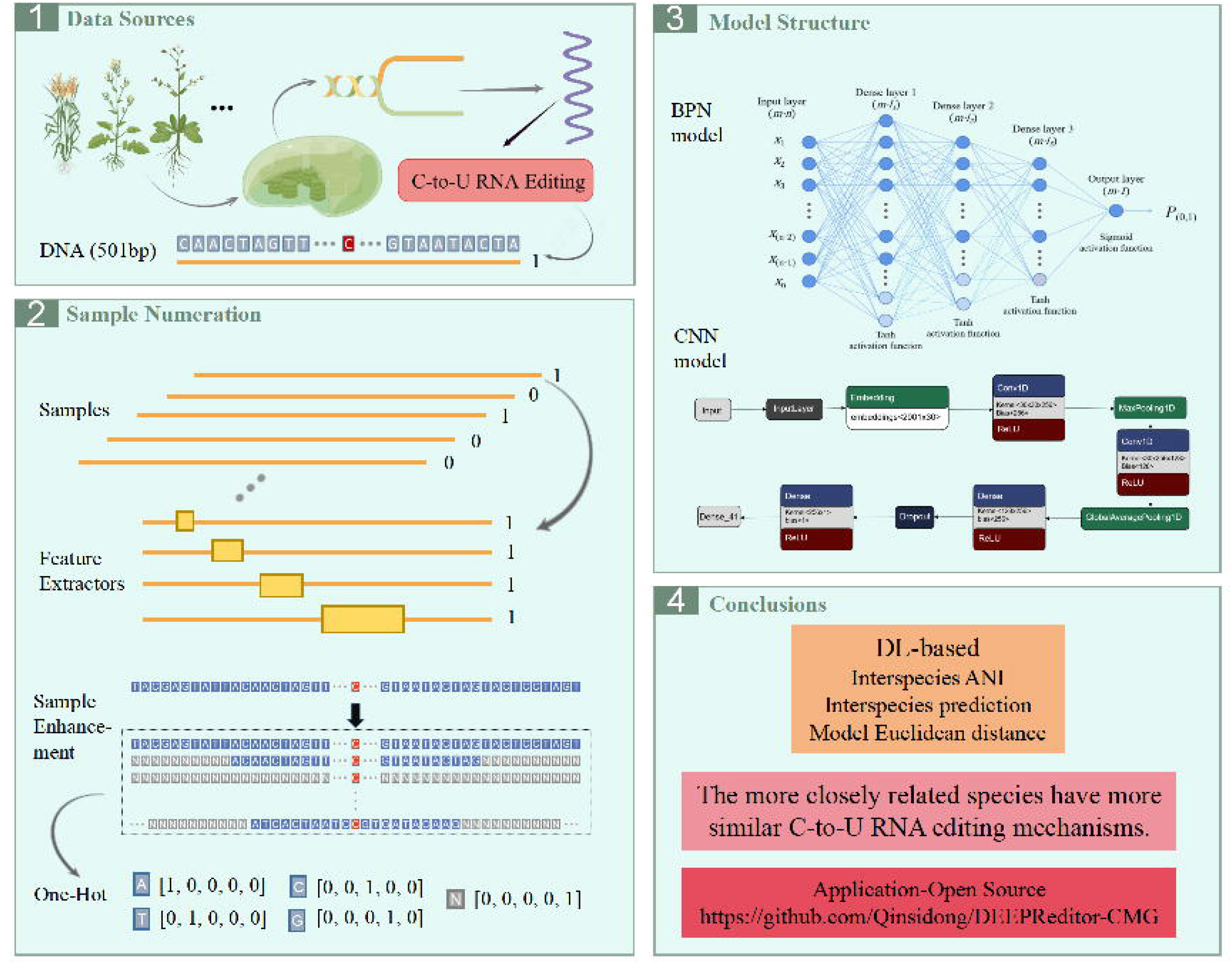

